# Recurrent circuits amplify corticofugal signals and drive feedforward inhibition in the inferior colliculus

**DOI:** 10.1101/2022.02.03.478995

**Authors:** Hannah M. Oberle, Alexander N. Ford, Jordyn E. Czarny, Meike M. Rogalla, Pierre F. Apostolides

## Abstract

The inferior colliculus (IC) is a midbrain hub critical for perceiving complex sounds such as speech. In addition to processing ascending inputs from most auditory brainstem nuclei, the IC receives descending inputs from auditory cortex that control IC neuron feature selectivity, plasticity, and certain forms of perceptual learning. Although corticofugal synapses primarily release the excitatory transmitter glutamate, many physiology studies show that auditory cortical activity has a net inhibitory effect on IC neuron spiking. Perplexingly, anatomy studies imply that corticofugal axons primarily target glutamatergic IC neurons while only sparsely innervating IC GABA neurons.

Corticofugal inhibition of the IC may thus occur largely independently of feedforward activation of local GABA neurons. We shed light on this paradox using *in vitro* electrophysiology in acute IC slices from fluorescent reporter mice of either sex. Using optogenetic stimulation of corticofugal axons, we find that excitation evoked with single light flashes is indeed stronger in presumptive glutamatergic neurons compared to GABAergic neurons. However, many IC GABA neurons fire tonically at rest, such that sparse and weak excitation suffices to significantly increase their spike rates.

Furthermore, a subset of glutamatergic IC neurons fire spikes during repetitive corticofugal activity, leading to polysynaptic excitation in IC GABA neurons owing to a dense intra-collicular connectivity. Consequently, recurrent excitation amplifies corticofugal activity, drives spikes in IC GABA neurons, and generates substantial local inhibition in the IC. Thus, descending signals engage intra-collicular inhibitory circuits despite apparent constraints of monosynaptic connectivity between auditory cortex and IC GABA neurons.

**Significance Statement:** Descending “corticofugal” projections are ubiquitous across mammalian sensory systems, and enable the neocortex to control subcortical activity in a predictive or feedback manner. Although corticofugal neurons are glutamatergic, neocortical activity often inhibits subcortical neuron spiking. How does an excitatory pathway generate inhibition? Here we study the corticofugal pathway from auditory cortex to inferior colliculus (IC), a midbrain hub important for complex sound perception. Surprisingly, cortico-collicular transmission was stronger onto IC glutamatergic compared to GABAergic neurons. However, corticofugal activity triggered spikes in IC glutamate neurons with local axons, thereby generating strong polysynaptic excitation and feed-forward spiking of GABAergic neurons. Our results thus reveal a novel mechanism that recruits local inhibition despite limited monosynaptic convergence onto inhibitory networks.

## Introduction

Feedback projections from the sensory neo-cortex to sub-cortical regions are ubiquitous in the mammalian brain. These corticofugal pathways enable “high-level” cortical computations to rapidly control the nature of ascending sensory signals, hypothetically supporting important “top-down” functions such as predictive coding, error propagation, or stream segregation (Briggs and Usrey, 2011; Stebbings et al., 2014; Usrey and Sherman, 2019; Asilador and Llano, 2020). Interestingly, corticofugal activity often has a net inhibitory effect upon spontaneous and sensory-evoked activity in sub-cortical circuits, consequently sharpening receptive fields and increasing the sparseness of neuronal representations along the ascending hierarchy (Syka and Popelár, 1984; Zhang et al., 1997; Bledsoe et al., 2003; Boyd et al., 2012; Crandall et al., 2015; Vila et al., 2019; Born et al., 2021; but see Kirchgessner et al., 2021). Because corticofugal projections originate almost exclusively from excitatory (glutamatergic) neurons in cortical layers 5 and 6 (Hattox and Nelson, 2007; Schofield, 2009; Slater et al., 2013; Asilador and Llano, 2020; Sherman and Usrey, 2021), the inhibitory consequences of corticofugal activity may occur via feedforward GABAergic interneurons. However, the cellular and circuit-level mechanisms that enable corticofugal activity to reliably generate sub-cortical inhibition are poorly understood; this knowledge gap persists owing to the difficulty of quantifying corticofugal transmission in identified excitatory and inhibitory neurons.

In the auditory system, the corticofugal projection from auditory cortex to the inferior colliculus (IC) is of particular importance as the IC relays the majority of ascending acoustic signals destined for forebrain circuits. Accordingly, corticofugal activity powerfully shapes how IC neurons respond to diverse sound features, often via inhibitory interactions. Stimulating the auditory cortex dampens or completely suppresses IC acoustic responses (Syka and Popelár, 1984; Bledsoe et al., 2003; Vila et al., 2019; Blackwell et al., 2020), and intracellular data show that auditory cortex stimulation drives synaptic inhibition in individual IC neurons (Mitani et al., 1983; Qi et al., 2020). Conversely, silencing auditory cortex often potentiates spontaneous and sound-evoked activity in the IC (Nwabueze-Ogbo et al., 2002; Popelár et al., 2003; Popelář et al., 2016; but see Zhang and Suga, 1997), indicating that ongoing auditory cortical activity can have a net inhibitory effect on the moment-to-moment excitability of IC neurons. However, anatomical data show that IC GABA neurons receive surprisingly few synapses from auditory cortex; most corticofugal axons instead target glutamatergic IC neurons (Nakamoto et al., 2013; Chen et al., 2018). Thus, rather than recruiting *local* inhibitory neurons to sharpen IC neuron tuning, auditory cortex may instead exert inhibitory control through long-range inhibitory pathways such as the dorsal nucleus of the lateral lemniscus (Beneyto et al., 1998; Budinger et al., 2000), via local effects *upstream* of the IC (Kong et al., 2014), or through a newly discovered, monosynaptic GABAergic corticofugal projection (Bertero et al., 2021). Alternatively, circuit mechanisms beyond the orthodox feedforward inhibitory motif, whereby long-range excitation preferentially drives GABAergic interneurons (Pouille and Scanziani, 2001; Gabernet et al., 2005; Cruikshank et al., 2007; Boyd et al., 2012), may enable the auditory cortex to generate local inhibition in the IC.

We used patch-clamp electrophysiology and optogenetics in brain slices from fluorescent GABA reporter mice to study corticofugal transmission onto identified IC neurons. We focused on neurons in the dorso-medial “shell” IC because this sub-region receives the densest projection of corticofugal axons (Winer et al., 1998; Song et al., 2018; Oberle et al., 2022). Our data confirm anatomical predictions by showing that the bulk strength of corticofugal transmission is greater onto presumptive glutamate neurons compared to GABA neurons in the dorsal IC. However, dorsal IC GABA neurons are densely contacted by potent, intra-collicular synapses from local glutamate neurons. As such, repetitive corticofugal activity drives strong polysynaptic excitation in a subset of dorsal IC GABA neurons and generates local inhibition. Our results identify a novel mechanism for the neo-cortex to drive inhibition in a sub-cortical pathway. More broadly, the data show how recurrent activity in local circuits sign-inverts excitatory corticofugal signals.

## Materials and Methods

The experiments were approved by the University of Michigan’s IACUC and performed according to the NIH’s Guide for the Care and Use of Laboratory Animals. *In vitro* experiments were performed on 5-9 week old male or female offspring of VGAT-ires-cre x Ai14 fl/fl (Jackson Labs stock # 028862 and 007914, respectively) breeder pairs from our colony (Figures 2-10) or 5-8 week old male and female C57/Bl6J mice ordered from Jackson labs (Figure 11; stock # 000664). For *in vivo* experiments with wild-type mice (of either sex), we used 4-7 week old offspring of CBA/CaJ (stock # 000654) x C57Bl6/J matings (Figure 1A-E) or 7-10 week old C57/BL6J mice ordered from Jackson labs (Figure 1F-J). The raw data from some of these *in vivo* recordings were also included in a previous set of distinct analyses (Oberle et al., 2022).

**Figure 1:**
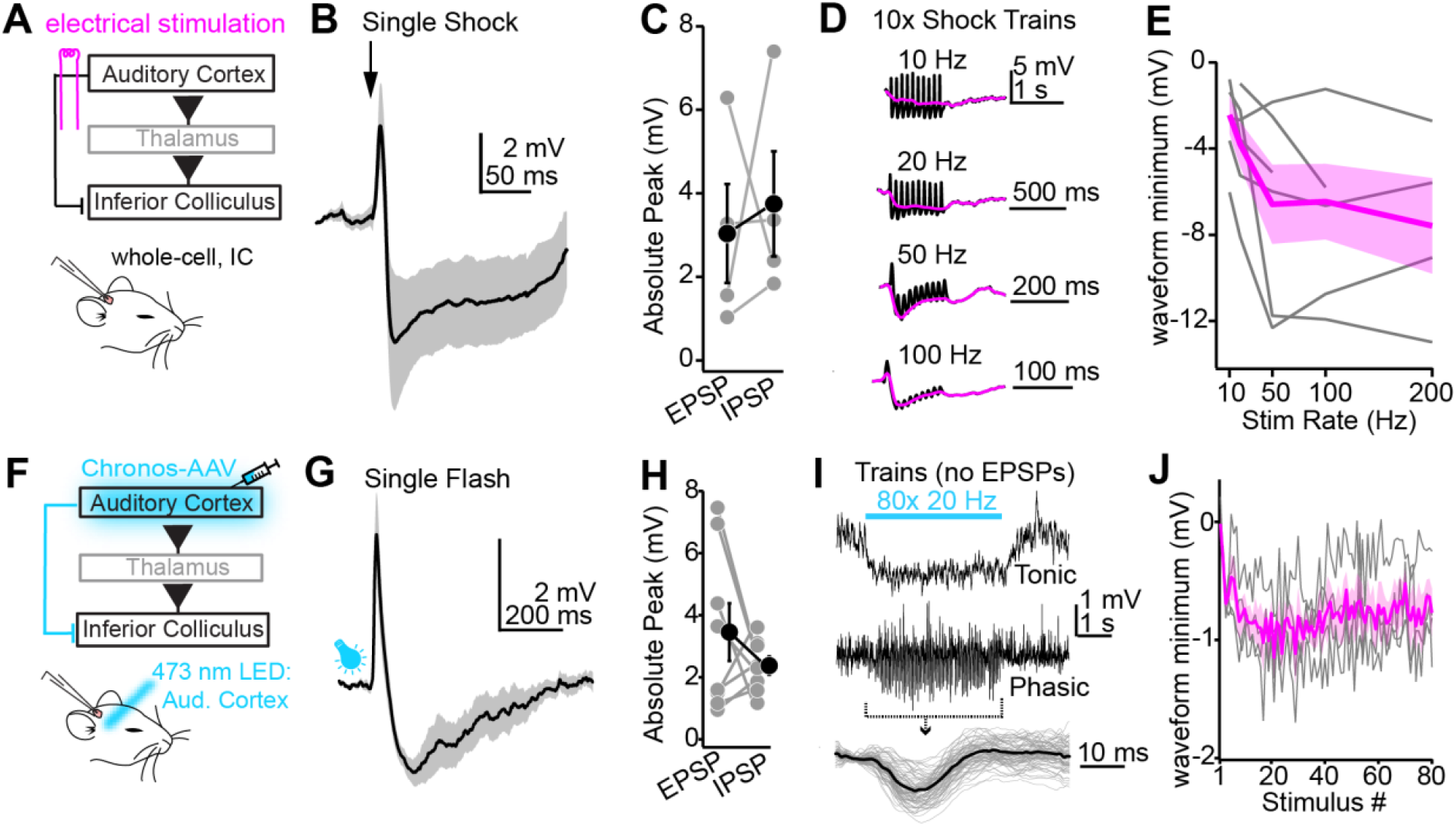
Auditory cortical stimulation generates IPSPs in IC neurons. **A)** Cartoon of experiment. Whole cell recordings are obtained from superficial IC neurons in urethane anesthetized mice; a bipolar electrode is placed in auditory cortex to activate corticofugal fibers. **B)** Single 20 µs shocks to the auditory cortex evoke a brief EPSP followed by a large and long-lasting IPSP. Trace is mean ± SEM from n = 4 neurons under these conditions **C)** Summary of EPSP and IPSP peak amplitude from the 4 cells averaged in panel B. Gray are individual data points, black is mean ± SEM. Lines connect individual experiments. **D)** Black: Example average membrane potential changes i during auditory cortex stimulation trains of different rates. Magenta is the mean tonic membrane potential, calculated by blanking phasic PSPs via linear interpolation and smoothing the waveform with a 50 ms sliding window. Examples shown are averages from n = 4 to 6 cells per condition, as not all stimulation rates were tested in each experiment. **E)** Summary of the peak tonic hyperpolarization during different auditory cortical train stimuli shown in panel D. Gray lines are individual cells; Magenta is mean ± SEM. **F)** Chronos is expressed in auditory cortex neurons via AAV injections; 2-4 weeks later, whole-cell recordings are obtained from superficial IC neurons. **G)** Example average traces showing EPSP-IPSP sequences following single light flashes delivered to the auditory cortex. Trace and shading are mean ± SEM from n = 8 neurons. **H)** Summary data from recordings as in panel G. Coloring is same as in panel C. **I)** In a subset of neurons, repetitive optogenetic stimulation (80 flashes at 20 Hz) generates no apparent EPSPs, but rather tonic or phasic hyperpolarizations of the membrane potential (upper and middle traces, respectively). Traces are averages of multiple trials from two different neurons. Lower panel: Individual phasic IPSPs from the recording in the middle panel are aligned (gray) and averaged (black). **J)** Summary quantifying membrane potential hyperpolarization throughout the 80x 20 Hz optogenetic train in n = 4 neurons without apparent corticofugal excitation. Gray traces are individual experiments, magenta is mean ± SEM.

**Figure 2:**
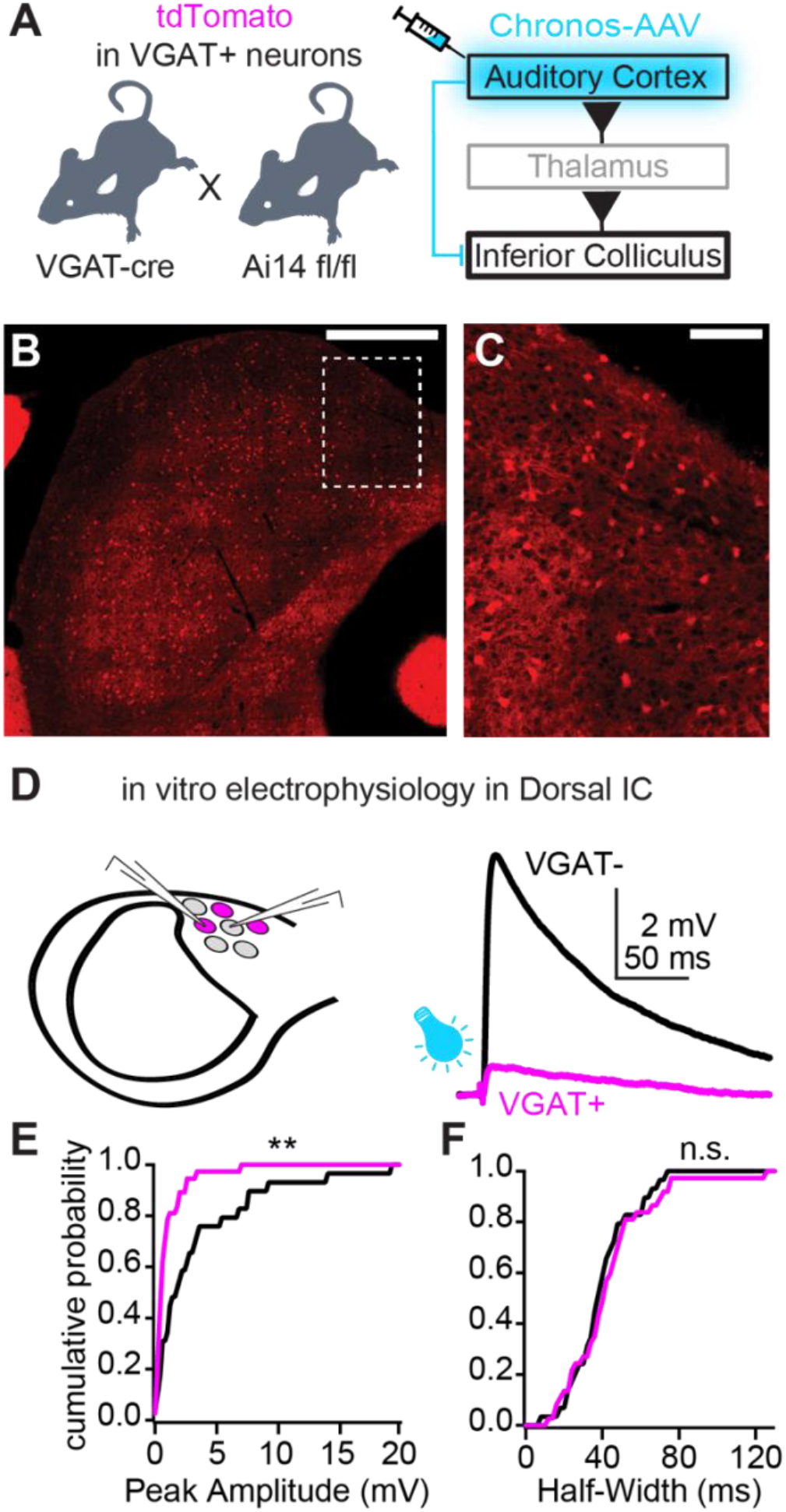
Differential strength of corticofugal EPSPs onto dorsal IC VGAT-and VGAT+ neurons. **A)** Diagram of experiment. The optogenetic activator Chronos is virally transduced into auditory cortex neurons of transgenic VGAT-cre x Ai14 mice. **B)** Tile scan of the IC in a VGAT-cre x Ai14 mouse. An area of interest is denoted by the dashed line and shown at higher magnification in panel C. Scale bar = 500 µm. **C)** Magnification of dashed rectangle in panel B. The micrograph was contrast enhanced to highlight tdTomato-positive presumptive GABA neurons and tdTomato-negative, presumptive glutamate neurons visible as dark “shadows”. Scale bar = 100 µm. **D)** 2-4 weeks following surgery, optogenetically evoked EPSPs are recorded in visually targeted tdTomato-positive and negative neurons. Right, example average EPSPs evoked by single light flashes in a VGAT-(black) and VGAT+ (magenta) dorsal IC neuron. **E,F)** Summary data showing cumulative probability distributions for EPSP peak amplitudes (E) and half-widths (F) in VGAT-and VGAT+ neurons. ** p < 0.01.

### Surgery for intracranial virus injections

Our virus injection protocol is described in detail in Oberle et al., 2022. Briefly, mice were anesthetized with 4-5% isoflurane in O2, mounted in a stereotaxic frame, and isoflurane was lowered to 1-2% while body temperature was maintained near 37-38° C using a heating blanket. Mice were administered 5 mg/kg carprofen as a pre-surgical analgesic, a small incision was made in the scalp, and 2% lidocaine was then applied to the wound margins. A 200-400 µm craniotomy was carefully drilled over the left auditory cortex (-2.75 mm from bregma, centered on the lateral ridge) or left IC (0.9 mm caudal and 1 mm lateral from lambda suture) to inject 100-200 nL of pAAV-Syn-Chronos-GFP virus (Addgene #59170-AAV1). At the end of the surgery, the craniotomy was filled with bone wax and the skin was sutured. Immediately following surgery, mice received an analgesic injection of buprenorphine (0.03 mg/kg, s.c.) and recovered on a heating pad before returning to their home cage. A post-operative dose of carprofen was administered ∼24 hours later.

### In vivo electrophysiology

Mice were deeply anesthetized with isoflurane, mounted in the stereotaxic frame, and craniotomies were carefully opened over the left IC and auditory cortex as described above and in Oberle et al. 2022. For optogenetic stimulation, we left the dura intact and implanted a cranial window over the auditory cortex. For electrical stimulation we made a small slit in the dura and the craniotomy was sealed with silicone elastomer. The IC craniotomy was plugged with silicone elastomer, a titanium headbar was implanted with dental cement, and the mouse was removed from the stereotax before being re-anesthetized with urethane (1.5 g/kg, i.p.) and head-fixed in a custom-made sound attenuation chamber. Body temperature during the recording session was maintained at 37-38° C with a custom designed, feedback-controlled heating blanket. Optogenetic stimulation was performed with a 0.5 NA, 400 µm core optic fiber (Thorlabs M45L02) coupled to a 470 nm LED (Thorlabs M470F3) positioned <1 mm away from the auditory cortex cranial window. LED power (∼25 mW peak power) was similar across experiments and flash duration (1-5 ms) was titrated to evoke consistent excitatory postsynaptic potentials (EPSPs) with low temporal jitter. For electrical stimulation experiments, a bipolar platinum-iridium electrode (FHC 30210) was carefully inserted ∼800 µm into auditory cortex at an angle roughly perpendicular to the cortical layers and shocks were delivered via a custom stimulus isolator. Stimulus intensity (2-5 mA) and duration (20 to 50 µs pulse-width) were titrated so as to evoke short-latency EPSPs with minimal transmission failures. Whole-cell current-clamp recordings were obtained ∼100-400 µm from the dura with pipettes containing (in mM): 115 K-Gluconate, 4 KCl, 0.1 EGTA, 10 HEPES, 14 Tris-Phosphocreatine, 4 Mg-ATP, 0.5 Tris-GTP, 4 NaCl, pH 7.2-7.3, 290 mOsm (open tip resistance: 5-10 MΩ).

### In vitro electrophysiology

All recordings were exclusively performed in the left IC hemisphere, which was ipsilateral to the auditory cortical injection site. 2-4 weeks following viral injections, mice were deeply anesthetized with isoflurane and 200-300 µm brain slices containing both IC hemispheres were prepared in warm (∼34° C) oxygenated ACSF containing (in mM): 119 NaCl, 25 NaHCO3, 3 KCl, 1.25 NaH2PO4, 15 glucose, 1 MgCl2, 1.3 CaCl2, 1 ascorbate, 3 pyruvate. A small cut was typically made in the lateral portion of the right cerebellum or right IC to visually identify the un-injected hemisphere. Slices were then incubated at 34° C in a holding chamber filled ACSF for 25-30 min and then stored at room temperature until use. During experiments, slices were mounted in a submersion chamber continuously perfused with oxygenated ACSF heated to 32-34° C (2-4 mL/min; chamber volume: ∼ 1 mL). Neurons in the dorso-medial shell IC were visualized via DIC using a Zeiss Axioskop FS 2 Plus microscope (40x objective; experiments of Figures 3-5 and 7-11) or via Dodt contrast using an Olympus BXW51 (63x objective; experiments of Figures 2-8 and 10-11). GABAergic neurons were identified from tdTomato fluorescence. Presumptive glutamatergic neurons were identified based on lack of fluorescence in VGAT-cre x Ai14 mice. Whole-cell current-clamp recordings were obtained using pipettes filled with the same K^+^ rich internal solution as employed for in vivo recordings (open tip resistance: 3-6 MΩ). In some experiments, 30 µM Alexa 488 or 0.1% biocytin were added to the internal solution to visualize neuronal morphology via online fluorescence or post-hoc histological reconstruction. Voltage clamp recordings in Figures 5 and 6 were obtained with the K^+^ rich internal solution while clamping somata between -65 and -70 mV. Recordings from Figure 11 were obtained with a Cs^+^ based solution containing (in mM): 110 Cesium Methanesulfonate, 10 QX-314-Bromide, 0.1 EGTA, 10 HEPES, 0.5 Tris-GTP, 4.5 MgATP, 5 TEA-Cl, 10 Tris-phosphocreatine. Experiments were generally conducted within 3-4 hours following slice preparation. For experiments in Figure 2, recordings from VGAT-and VGAT+ neurons were interleaved in most experiments to minimize potential effects arising from variability in opsin expression. Chronos-expressing axons were stimulated via 2-5 ms wide-field flashes of blue light delivered through the 63x objective (6.4 mW peak power). This light intensity suffices to saturate optogenetic activation of corticofugal axons under our conditions (See Figure 2E of Oberle et al., 2022). For experiments of Figure 6, 200 µM NMDA was diluted in ACSF from a stock solution, and delivered in the vicinity of the recorded cell via pulled glass pipettes and pressure application (5 psi, 100 ms). ACSF was applied using the same conditions during vehicle-only control recordings.

**Figure 3:**
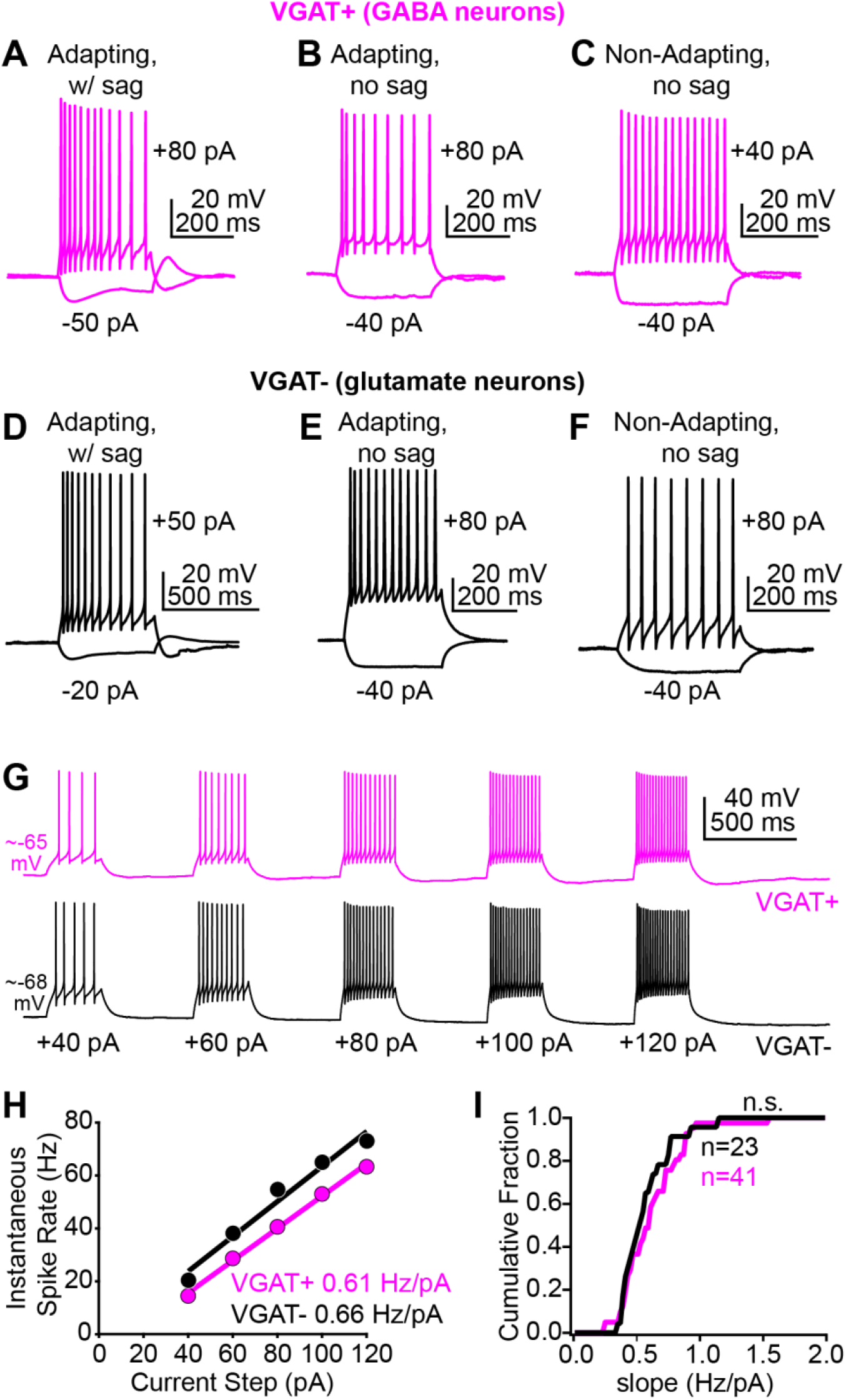
Dorsal *IC VGAT-and VGAT+ neurons have similar cellular properties.* **A-C)** Examples of distinct types of hyperpolarizing and depolarizing square pulse current steps in VGAT+ neurons encountered in the shell IC. **D-F)** Same as A-C, but for VGAT-neurons. Of note is the striking similarity in firing patterns and membrane properties of both neuron classes. **G)** Example spike responses to increasing square pulse current steps in a VGAT+ (magenta) and VGAT-(black) neuron. **H)** Instantaneous spike rate (y-axis) is plotted against current step amplitude (x-axis) for the two neurons of panel G. Lines are linear fits to the data, revealing similar slopes for the spike rate increases for the two neurons. **I)** Summary data showing a similar cumulative probability distribution of FI curve slopes for n = 41 VGAT+ neurons from N = 14 mice and n = 23 VGAT-neurons recorded in N = 11 mice (p = 0.5, Kolmogorov-Smirnov test).

**Figure 4:**
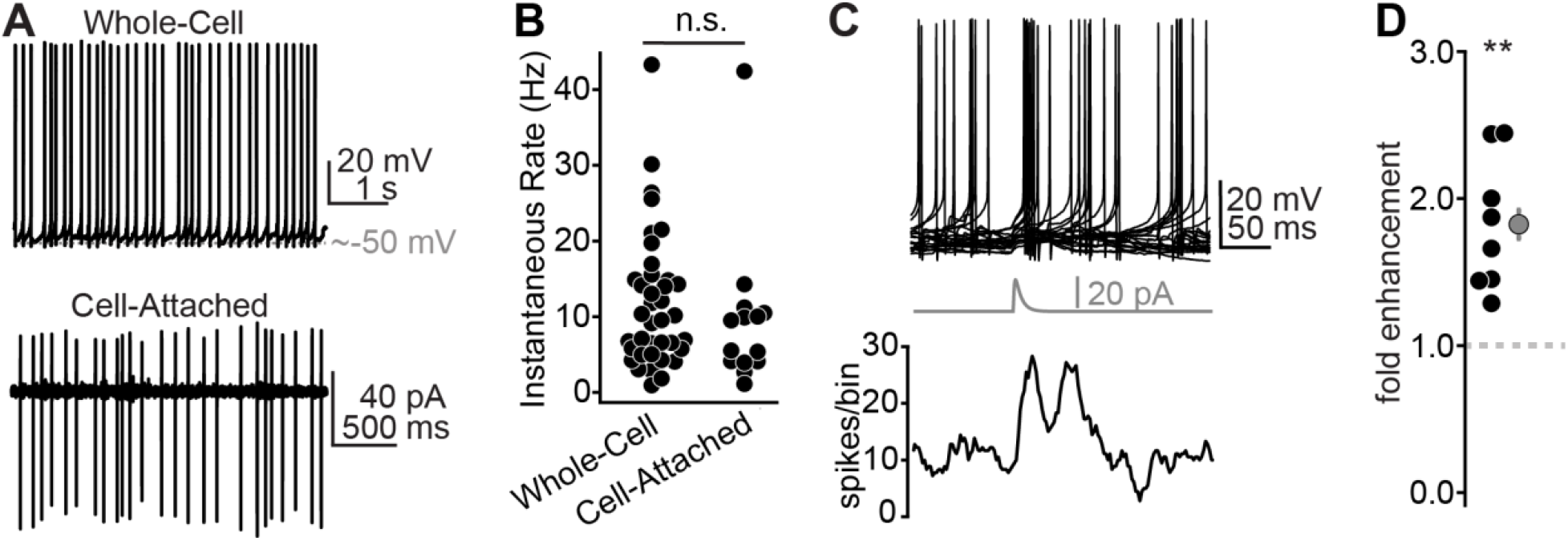
A subset of GABA neurons fire tonically, thus requiring only small currents to increase baseline firing rates. **A)** Example traces from tonic firing IC GABA neurons recorded in a VGAT-cre x Ai14 mouse in whole-cell or cell-attached modes (upper and lower panels, respectively). **B)** Summary data of tonic firing rates in IC GABA neurons recorded in the two configurations shown in panel A. **C)** Top: Example overlaid traces from a tonically firing IC GABA neuron. Middle: An EPSC-like current waveform (20 pA peak amplitude) increases spike probability in the ∼50 ms following current injection, as exemplified in the spike raster (lower panel). **D)** Summary data from n = 8 neurons as in panel C, showing that small EPSC waveforms (10-20 pA) nearly double the spike probability of tonically firing GABA neurons. ** p < 0.01.

**Figure 5:**
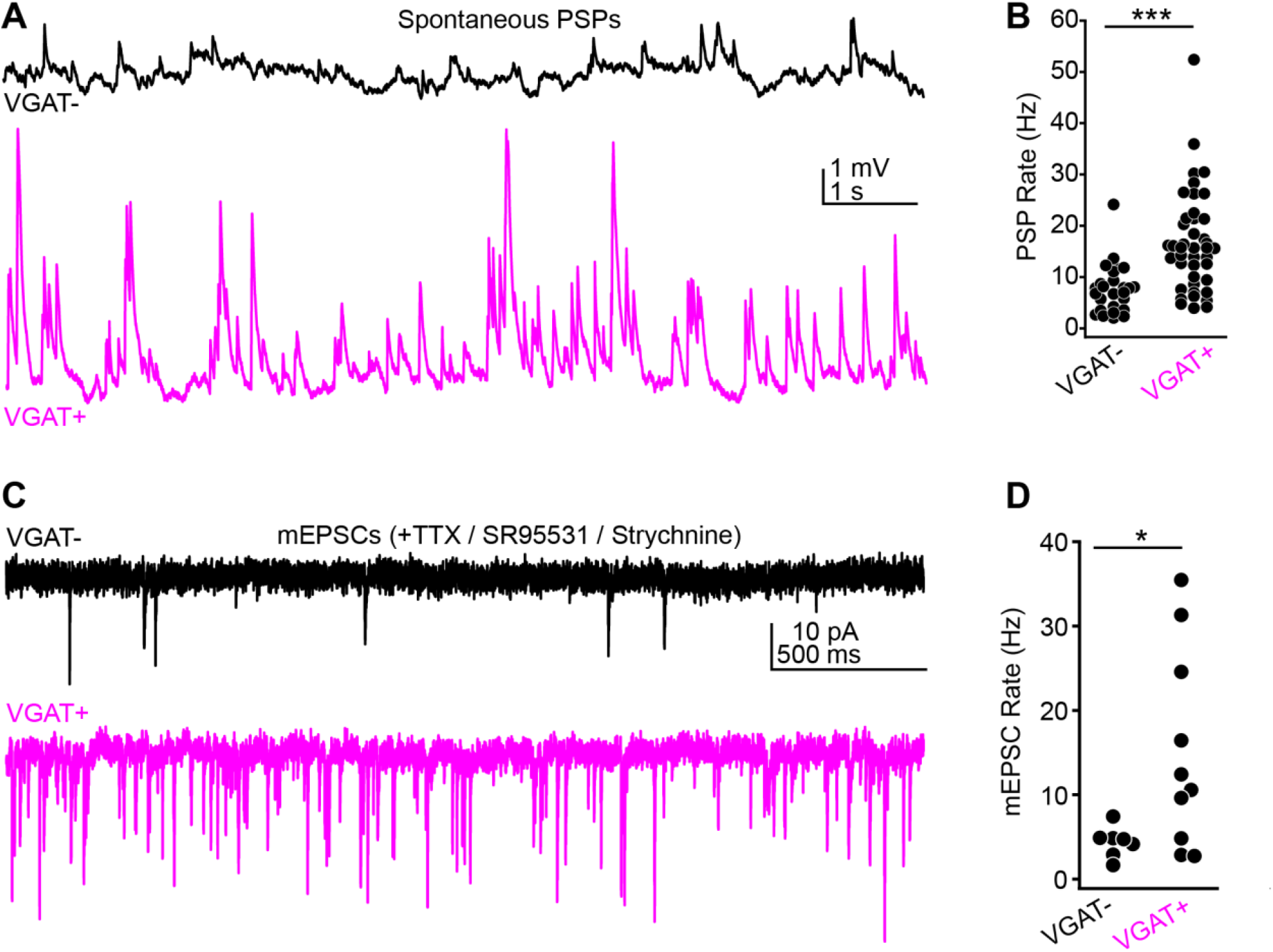
Differential convergence of excitatory synapses onto dorsal IC VGAT-and VGAT+ neurons. **A)** Examples of spontaneous synaptic activity in single VGAT-(black) or VGAT+ (magenta) dorsal IC neurons. Scale bars apply to both panels. **B)** Summary of instantaneous PSP rates for the two neuron classes. **C)** Example mEPSCs recorded in VGAT-and VGAT+ neurons. Color scheme is same as panel A. **D)** Summary of mEPSC rates across the two neuron classes. Similar to results with spontaneous PSPs, the instantaneous mEPSC rate was significantly higher in VGAT+ compared to VGAT-neurons. * p < 0.05, *** p < 0.001.

**Figure 6:**
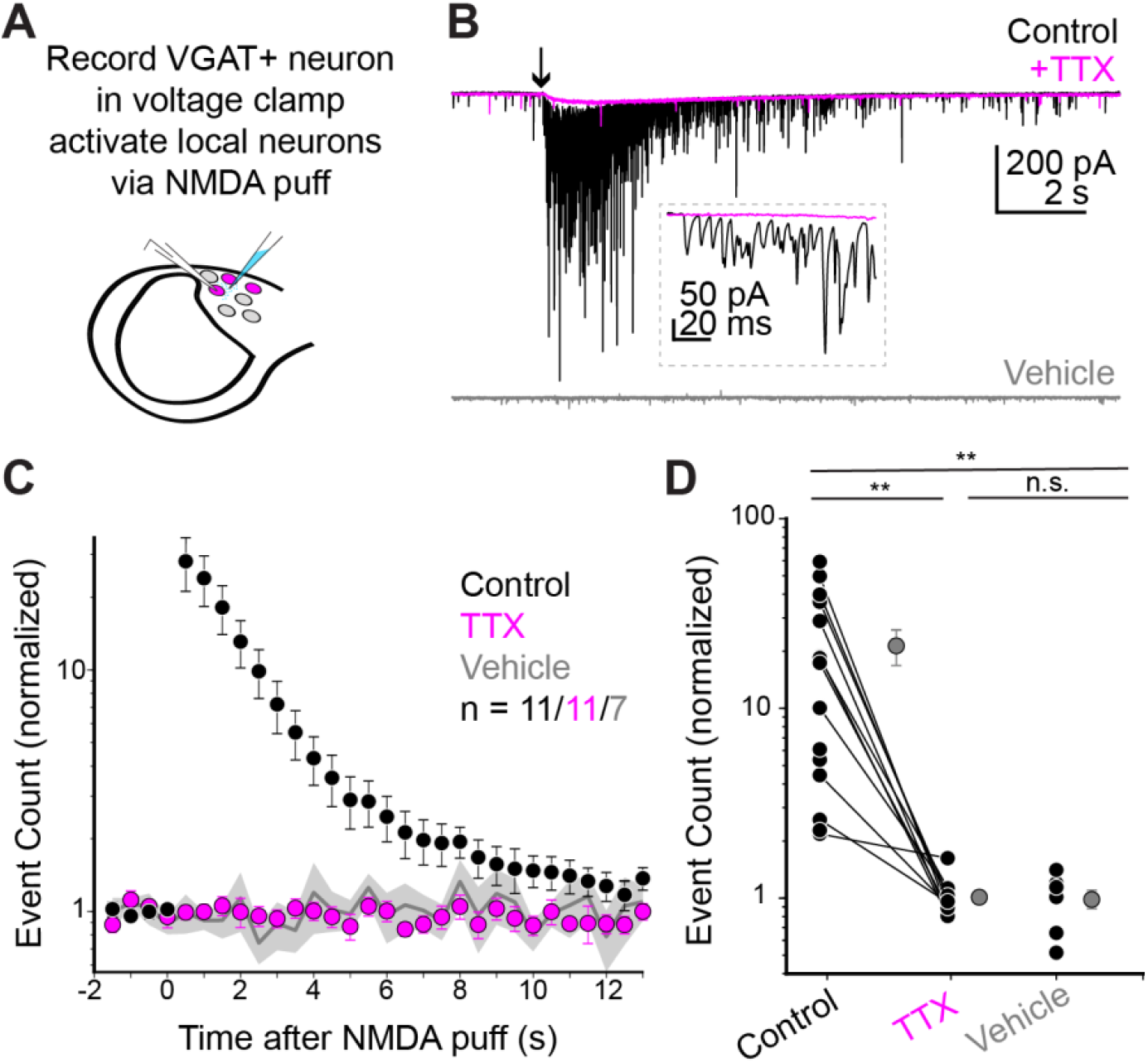
Dorsal IC VGAT+ neurons receive powerful intra-collicular excitation. **A)** Diagram of experiment. Whole-cell voltage-clamp recordings are obtained from tdTomato-positive VGAT+ neurons in the dorso-medial shell of the IC. NMDA (200 µM) was puff applied in the vicinity of the neuron. **B)** Example responses in control conditions (black) and following bath application of TTX (1 µM; magenta). Arrow denotes time of NMDA puff. Of note is the drastic increase in sEPSC rate in control that is blocked by TTX, indicating spike-driven release. Lower trace: Vehicle-only (ACSF) control shows no increase in sEPSC rate. **C)** Summary data of NMDA evoked event count increases in control and TTX. Bin-width is 500 ms, data are normalized to the event counts in a 2s baseline prior to NMDA puff and plotted on a log-scale. Vehicle puff summary data is plotted in background (gray line). Of note are the similar event rates in vehicle and TTX recordings. Error bars and shaded region are ± SEM. **D)** Summary data of the mean normalized event count in a 500 ms bin for the 2 seconds following the NMDA puff. Black and gray points represent individual experiments and mean ± SEM, respectively. ** p < 0.01.

**Figure 7:**
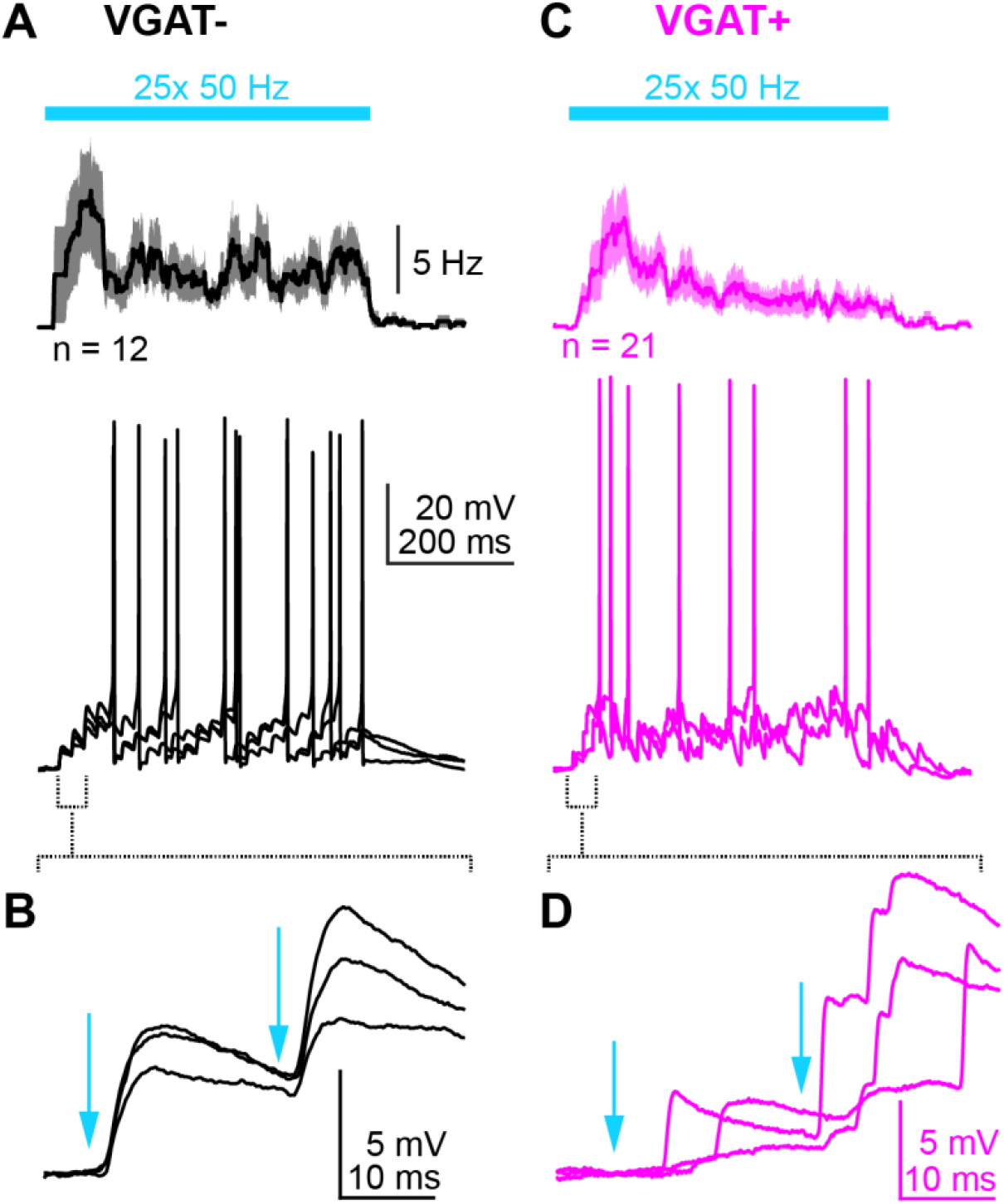
Repetitive corticofugal activity drives spikes in VGAT-and VGAT+ neurons. **A)** Upper panel: Average spike rate histogram for n = 12 VGAT-neurons during corticofugal train stimulation (n = 7 whole-cell recordings from N = 6 mice; n = 5 cell-attached attached recordings from N = 3 mice). Lower panel shows three overlaid trials from an example experiment. **B)** EPSPs following the first two light flashes are shown at a faster timebase. **C)** Same as A, but for VGAT+ neurons (n = 21 whole-cell recordings from N = 15 mice). **D)** Same as B, but for the example VGAT+ recording. Of note, the jittered onset and barrage-like nature of the synaptic events following the light flashes.

**Figure 8:**
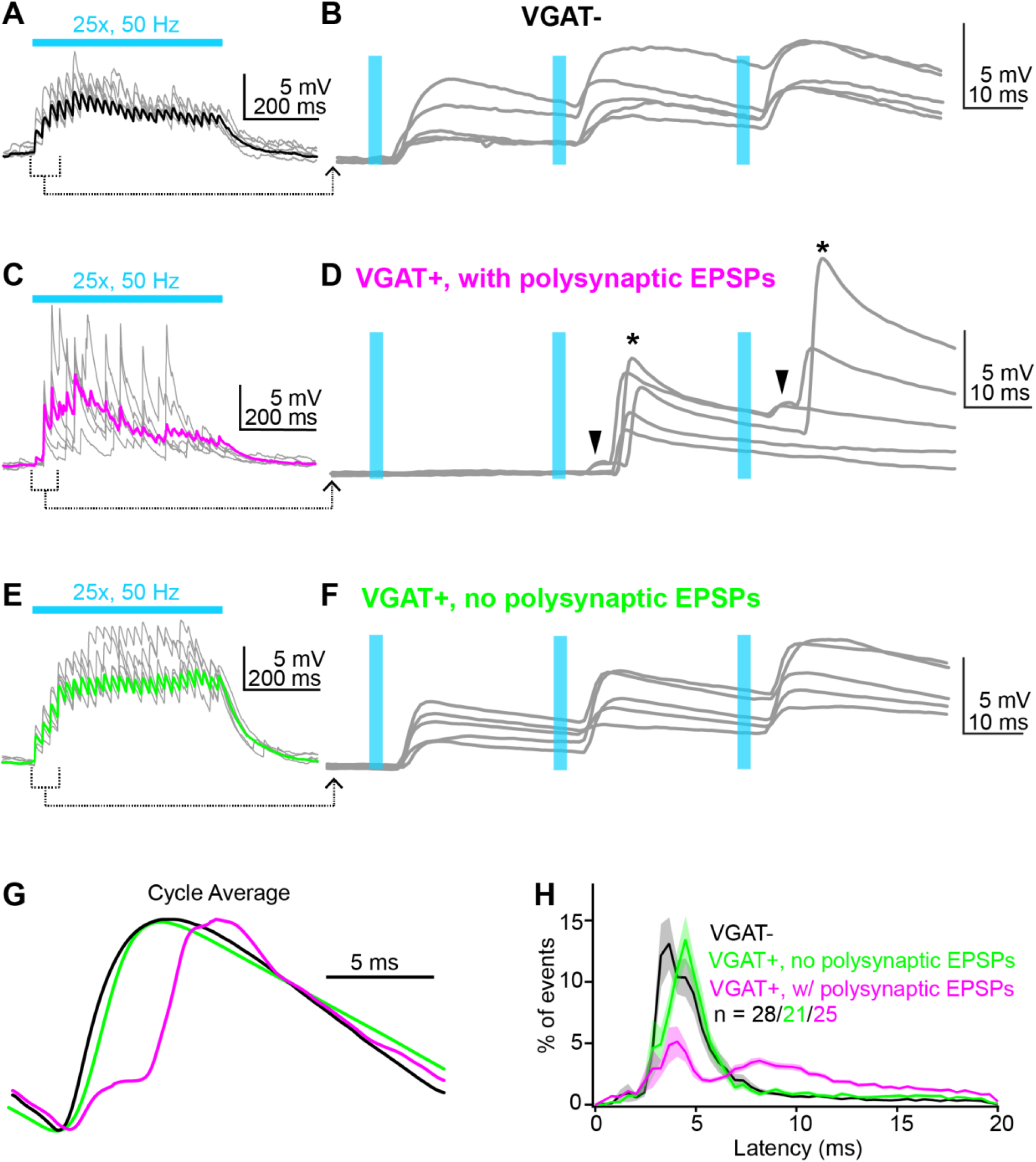
Repetitive corticofugal activity triggers polysynaptic EPSPs in a subset of VGAT+ IC neurons. **A)** Example EPSPs in a VGAT-neuron during train stimulation of corticofugal axons. Gray traces are individual trials, black is average of multiple trials. **B)** The first through third stimuli from panel A are shown at faster timebase. Blue lines denote light flashes. Of note is the short latency and low onset jitter. **C,D)** Same layout as in A-B, but for a VGAT+ neuron showing polysynaptic activity during corticofugal stimulation. Of note is the long latency and jittered onset of the large EPSPs, a hallmark of polysynaptic origin (asterisks). **E,F)** In another VGAT+ neuron, repetitive corticofugal activity generates primarily monosynaptic EPSPs similar to VGAT-neurons. Layout is same as A-D. **G)** The membrane voltage following each stimulus in the 50 Hz trains is averaged and peak normalized. Black, magenta, and green traces are data from the example VGAT-and VGAT+ neurons shown above, respectively. Of note is the dual component EPSP in the VGAT+ neuron with long-latency EPSPs, suggesting a monosynaptic corticofugal input followed by a much larger recurrent EPSP from the local circuit. **H)** Summary of normalized EPSP latency distributions for n = 28 VGAT-neurons from N = 14 mice (black), n = 25 VGAT+ neurons from N = 15 mice with polysynaptic EPSPs (magenta), and n = 21 VGAT+ neurons from N = 14 mice without polysynaptic EPSPs (green). Shading is ± SEM.

### Data Acquisition

Data were acquired using a Molecular Devices Multiclamp 700B or AM Systems model 2400 patch-clamp amplifier, online filtered at 2-10 kHz, and digitized at 50 kHz with a National Instruments PCI-6343 card + BNC2090A interface controlled by Matlab based acquisition software (Wavesurfer). As shown in Figure 4, many GABA neurons fired tonically *in vitro*. Consequently, negative bias current (-5 to -200 pA) was often injected to hyperpolarize neurons (typically between -60 and -80 mV) and prevent spontaneous or optogenetically triggered spiking when recording EPSPs in VGAT+ neurons. As such, the majority of spontaneous firing datasets of Figure 4 were recorded within the first few minutes of commencing whole-cell configuration, prior to hyperpolarizing the neurons to record optically evoked auditory cortical EPSPs.

### Data analysis

All experiments were analyzed using custom scripts written in Matlab. For experiments in Figure 1, stimulus artifacts and action potentials evoked by auditory cortex stimulation were blanked via linear interpolation. For analyses of tonic hyperpolarization in Figures 1D-E, phasic EPSPs were similarly blanked via linear interpolation and the waveform was smoothed with a 50 ms sliding window. Analyses in Figures 1, 2, 10, and 11E were run on averages of multiple trials (typically >10 trials per condition) after baseline subtraction and lowpass filtering at 1 kHz. EPSP peak amplitudes were calculated by averaging data points ± 0.1 ms around the local maximum following optogenetic stimulation. Halfwidths were defined as the full width at half-maximum of the peak. Voltage integrals in Figure 10 were calculated using the Matlab functions trapz() or cumtrapz(). Event detection analyses in whole-cell recordings from Figures 4C-D, 5A-B, 7-9, and 11A-D were conducted similar to previous approaches (Morales et al., 1985; Cocatre-Zilgien and Delcomyn, 1990; Ankri et al., 1994): We differentiated the membrane voltage or current traces, smoothed the data (1-10 ms sliding window), and used open source Matlab routines (peakfinder or crossdet; Matlab file exchange #25500 and 47266, respectively) to identify synaptic events and action potentials based on a rate of rise threshold crossing. For analysis of action potentials in cell-attached recordings (Figure 4A,B; some experiments of Figure 7), event detection routines were run on bandpass filtered (300-1000 Hz) current-or voltage-clamp data rather than differentiated traces. Miniature EPSCs in Figures 5C,D were detected using the template matching algorithm in AxographX (Clements and Bekkers, 1997), whereas sEPSC detection in Figure 6 was performed using a deconvolution approach similar to Pernía-Andrade et al., 2012. Prior to deconvolution, the contribution of postsynaptic NMDA currents was removed from each trace by digitally subtracting a low-pass filtered (50 Hz) average of NMDA responses recorded in TTX. Synaptic current waveforms (Figure 4C,D) were generated in Wavemetrics Igor Pro using the Neuromatic package (Rothman and Silver, 2018).

### Statistics

In the text, n and N refer to the number of recorded neurons and mice, respectively. Sample sizes were not explicitly pre-determined prior to data collection. However, the number of cells and mice in these experiments are in accordance with commonly accepted standards in the field. Data were tested for normality using a Lilliefors test prior to statistical comparisons and for equal variances using Levene’s test where appropriate. Parametric tests were used for normally distributed data, while non-parametric tests were used if one or more of the datasets deviated from normal. Alpha is corrected for multiple comparisons in post-hoc significance tests following ANOVA. Repeated measures ANOVA were run using Geisser-Greenhouse correction. Statistics were run in Matlab or Graphpad Prism 9. Pearson’s coefficient and associated bootstraps were calculated in Matlab with the functions corr() and bootstrp(), respectively.

## Results

### Cortical stimulation inhibits superficial IC neurons in vivo

Previous studies show that stimulating the auditory cortex profoundly reduces sound-evoked spiking in IC neurons (Syka and Popelár, 1984; Bledsoe et al., 2003). In our recent *in vivo* whole-cell recordings from cortico-recipient neurons in the superficial IC layers (see also Oberle et al., 2022 for complementary analyses of these datasets), we noticed that auditory cortex stimulation triggered putative feedforward inhibitory postsynaptic potentials (IPSPs) which might account for these prior observations. When stimulating the auditory cortex with a bipolar electrode (Figure 1A), single electric shocks (20 µs pulse-width) drove excitatory postsynaptic potentials (EPSPs) followed by large inhibitory postsynaptic potentials (IPSPs) in 4/7 cells tested (Figure 1B,C. EPSP and IPSP peak amplitudes: 3.04 ± 1.18 and 3.75 ± 1.26 mV, respectively; n = 4 cells from N = 2 mice). In the other three cells we only observed EPSPs after single shocks, although IPSPs could be elicited in 2/3 of these neurons by increasing the stimulus duration from 20 to 50 µs. IPSPs often summated during repetitive train stimulation (10 shocks, 50 µs pulse-width, delivered between 10-200 Hz), such that train evoked EPSPs typically initiated atop a tonic membrane potential hyperpolarization (Figure 1D). Interestingly, cortically mediated hyperpolarizations could be evoked in all neurons tested using at least one stimulus train condition (n = 7 neurons from N = 4 mice), with faster stimulation rates generally resulting in larger hyperpolarizations (Figure 1E).

We noticed similar corticofugal-driven IPSPs in a subset of recordings using optogenetic stimulation of auditory cortex neurons expressing the light gated opsin Chronos (Figure 1F). In n = 21 superficial IC neurons receiving putatively monosynaptic corticofugal excitation, EPSP/IPSP sequences were observable in n = 8 neurons from N = 4 mice when stimulating auditory cortex neurons with single light flashes (Figure 1G,H; EPSP and IPSP peak amplitudes: 3.45 ± 0.93 and 2.38 ± 0.29 mV, respectively). Interestingly, inhibition was also seen in 5/17 IC neurons where cortical stimulation did not evoke EPSPs. In 4 neurons, repetitive auditory cortical activity (80 light flashes delivered at 20 Hz) caused a small tonic or phasic hyperpolarization of the membrane potential (Figure 1I,J; n = 2 neurons in each condition), whereas in 1 neuron tested, single light flashes generated a single, long-latency IPSP (data not shown). Altogether, these data indicate that in addition to the monosynaptic depolarizations mediated by glutamatergic corticofugal projections, descending activity also hyperpolarizes IC neurons, presumably via polysynaptic inhibition. Finally, IPSPs appeared more common with electrical compared to optogenetic stimulation. However this observation is likely incidental, as electrical stimulation likely causes stronger and/or more synchronous activation of corticofugal neurons to favorably recruit feedforward inhibition. Moreover, the absence of IPSPs in some recordings may simply reflect the fact that the degree of corticofugal driven inhibition depends on stimulation patterns which can be unique to the particular IC neurons under study (Vila et al., 2019).

### Corticofugal EPSPs are larger in VGAT-compared to VGAT+ IC neurons

The hyperpolarization of IC neurons observed during auditory cortex stimulation is intriguing, as anatomy data suggest that auditory cortico-collicular axons preferentially synapse onto excitatory glutamatergic IC neurons; far fewer descending synapses target GABA neurons in the IC (Nakamoto et al., 2013; Chen et al., 2018). We thus wondered if the functional strength of auditory corticofugal synapses differed between glutamatergic and GABAergic IC neurons in a manner consistent with anatomical, or rather, physiological data (e.g., Figure 1). To this end we expressed the red fluorescent protein tdTomato specifically in GABAergic neurons by crossing VGAT-ires-cre and Ai14 fl/fl mouse lines. We then we used stereotaxic injections of a pan-neuronal adeno-associated virus to broadly transduce the optogenetic activator Chronos in auditory cortex neurons of these mice. 2-4 weeks later, we prepared acute brain slices for whole-cell patch-clamp electrophysiology (Figure 2A). We targeted either tdTomato-positive (VGAT+) or -negative (VGAT-) somata in the dorso-medial shell IC to record from presumptive GABAergic or glutamatergic neurons, respectively (Figure 2B-D). Single flashes of blue light (2-5 ms duration, 6.4 mW) delivered through a 63x microscope objective activated Chronos-expressing auditory cortico-collicular axons and generated EPSPs that were >3-fold larger in VGAT-compared to VGAT+ neurons (Figure 2D,E. Median amplitude: 1.86 vs. 0.58 mV, n = 29 VGAT-neurons from N = 9 mice and n = 37 VGAT+ neurons from N = 10 mice, respectively. p = 0.0024, Kolmogorov-Smirnov test). By contrast, EPSP halfwidths were similar across the two neuron populations (Figure 2F; 37.5 vs. 41.0 ms, p = 0.7223, Kolmogorov-Smirnov test), implying that amplitude differences are unlikely due to greater recruitment of feedforward inhibition onto VGAT+ neurons. Additionally, this result cannot be accounted for by differences in driving force or input resistances across cell-types: Neurons were held at similar average membrane potentials during these recordings (-64.7 ± 1.2 mV vs. -63.6 ± 1.2 mV for VGAT-and VGAT+ neurons respectively, p = 0.52, unpaired t-test), and the median input resistance of VGAT-neurons was marginally, but significantly lower than that of VGAT+ neurons (320.9 vs. 427.5 MΩ, p = 0.04, Kolmogorov-Smirnov test). Moreover, EPSPs had similar short latency onsets in the two populations (3.4 vs 3.5 ms for VGAT-and VGAT+ neurons, p = 0.79, Kolmogorov-Smirnov test), suggesting that EPSPs predominantly reflect monosynaptic corticofugal inputs. In agreement with prior electron microscopy data showing that corticofugal synapses primarily target non-GABAergic structures in the IC, VGAT-neurons trended towards a higher probability of exhibiting EPSPs following light flashes compared to VGAT+ neurons (30/33, or 91% compared to 38/52, or 73% for VGAT-and VGAT+ neurons, respectively. p = 0.08, χ^2^ test with Yates correction). Altogether our data imply a specific connectivity logic to the auditory cortico-collicular pathway, whereby corticofugal transmission generates significantly larger EPSPs in VGAT-compared to VGAT+ neurons.

### A subset of dorsal IC VGAT+ neurons tonically fire action potentials

As expected from prior studies (Ono et al., 2005; Naumov et al., 2019), dorsal IC VGAT-and VGAT+ neurons had a range of overlapping electrical properties and could not be unambiguously distinguished by spike patterns (Figure 3A-F) or the slope of firing curves in response to current steps (Figure 3G-I). Aside from the moderate group-level difference in input resistance (see text above), VGAT-and VGAT+ neurons could thus not be easily distinguished on a case-by-case basis solely from membrane responses. However, we observed that VGAT+ neurons were significantly more likely to tonically fire spikes (38/72 VGAT+ neurons vs 6/37 VGAT-neurons showed spontaneous action potentials at rest, χ^2^ = 13.572, p < 0.001), such that recording sub-threshold EPSPs in VGAT+ neurons often necessitated injecting negative bias current to prevent tonic firing. These results were not an artifact of whole-cell dialysis, as VGAT+ neurons similarly fired spontaneously when recorded in “loose-patch” extracellular configuration (Figure 4A,B. Median spontaneous firing rates: 9.9 vs 7.5 Hz for whole-cell and loose-patch recordings, respectively. p = 0.15, rank-sum test; n=38 whole-cell recordings from N = 18 mice and n = 14 loose-patch recordings from N = 4 mice). Moreover, similar tonic firing was also reported in a subset of GABAergic IC neurons (Silveira et al., 2020). Consequently, we hypothesized that even sparse synaptic excitation might suffice to significantly increase the firing rate of tonically firing IC VGAT+ neurons. Indeed, injecting small excitatory postsynaptic current (EPSC)-like waveforms (10-20 pA peak amplitude) at the soma of tonically firing VGAT+ neurons significantly increased spike rates 1.82 ± 0.16 fold relative to a pre-current injection baseline period (Figure 4C,D; n = 8 neurons from N = 2 mice, p = 0.0012, one-sample t-test). Thus, a dense convergence of descending axons may not be an explicit pre-requisite to increase firing rates of some IC VGAT+ neurons.

### Dorsal IC VGAT+ neurons receive powerful intra-collicular excitation

In addition to differences in tonic firing, spontaneous PSP rates recorded at hyperpolarized membrane potentials were significantly higher in VGAT+ compared to VGAT-neurons (Figure 5A-B; median PSP rate 15.9 vs. 7.4 Hz, n = 43 VGAT+ neurons from N = 10 mice and n = 26 VGAT-neurons from N = 9 mice, respectively. p < 0.001, Kolmogorov-Smirnov test). Qualitatively similar results were obtained when measuring the rate of glutamatergic miniature excitatory postsynaptic currents (mEPSCs) isolated with SR95531 (5-10 µM), strychnine (500 nM), and tetrodotoxin (TTX; 1 µM) in voltage-clamp recordings (Figure 5C,D; 15.1 ± 3.7 vs 4.4 ± 0.7 Hz, n = 10 VGAT+ neurons from N = 4 mice and n = 7 VGAT-neurons from N = 4 mice, respectively. p = 0.0127, Welch’s t-test). By contrast, median mEPSC amplitudes were similar between the two groups (-12.8 vs. -10.3 pA, p=0.47 rank-sum test). These results indicate that the smaller amplitude of corticofugal EPSPs (Figure 2) does not reflect a paucity of excitatory synapses onto VGAT+ neurons, unusually small quantal size, or a preferential severing of VGAT+ neuron dendrites during slice preparation. Rather, the data corroborate anatomy studies suggesting that VGAT+ neurons receive a substantial amount of excitatory synapses from intra-collicular circuits (Ito and Oliver, 2014; Beebe et al., 2016; Schofield and Beebe, 2019), yet are contacted far less by descending inputs than VGAT-neurons.

The high rates of spontaneous synaptic excitation in VGAT+ neurons might arise from neighboring, glutamatergic IC neurons with local axon collaterals (Smith, 1992; Ito and Oliver, 2014). If true, spikes in neighboring neurons should increase the rate of excitatory synaptic events in VGAT+ neurons. We tested this hypothesis by voltage clamping somata of dorsal IC VGAT+ neurons to prevent spiking, and driving action potentials in neighboring presynaptic neurons via pressure application of NMDA in the vicinity of the recorded cell (Figure 6A). Local puffs of 200 µM NMDA (100 ms, 5 psi) rapidly and dramatically increased spontaneous EPSC (sEPSC) activity, causing a ∼20 fold peak increase in sEPSC rates above baseline during the 2 seconds following the puff (Figure 6C,D; n = 16 neurons from N = 13 mice; mean peak event rate increase ± SEM : 21.37 ± 4.67 fold above baseline). Importantly, the evoked increase in sEPSC was not observed during vehicle-only puffs (n = 7 neurons from N = 3 mice; mean peak event rate increase ± SEM: 0.99 ± 0.12), and it was abolished by bath application of 1 µM TTX (n = 11 neurons from N = 9 mice; mean normalized peak event rate ± SEM: 1.01 ± 0.07), resulting in a significantly greater peak rate of sEPSCs following NMDA application in control conditions (Welch ANOVA W(2.00, 16.15) = 9.13, p = 0.0022, Dunnett’s post-hoc tests: control vs. TTX p = 0.0017, control vs. vehicle p = 0.0017, TTX vs. vehicle p = 0.9985). NMDA puffs in TTX caused only a small (-23.82 ± 4.19 pA) inward current which presumably reflects agonist binding to NMDA receptors on the voltage-clamped, postsynaptic neuron. Thus, NMDA puffs increase sEPSC rates by driving spikes in local IC neurons, rather than via a direct depolarization of nerve terminals via presynaptic NMDA receptors (Glitsch and Marty, 1999; Rossi et al., 2012). Altogether, our results support the hypothesis that VGAT+ neurons receive substantial excitatory input from local, presynaptic glutamatergic IC neurons.

### Repetitive corticofugal activity drives spikes in VGAT-neurons, leading to polysynaptic EPSPs in VGAT+ neurons

The experiments in Figures 5 and 6 indicate that glutamatergic neurons of the dorsal IC synapse onto neighboring VGAT+ neurons. If descending synapses onto glutamatergic IC neurons were powerful enough to drive spikes, corticofugal activity could trigger feedforward excitation in GABAergic IC neurons. In this context, the IC’s recurrent excitatory circuitry would amplify auditory corticofugal signals and might subsequently drive spiking in GABAergic dorsal IC neurons. As a first test of this hypothesis, we asked if repetitively stimulating Chronos-expressing corticofugal axons (25 flashes at 50 Hz) depolarizes VGAT-and VGAT+ neurons beyond spike threshold; these stimulation parameters are within the physiological range of sound-evoked firing rates in corticofugal neurons (Williamson and Polley, 2019). In agreement, train stimulation triggered spikes in n = 7/29 (24%) of VGAT-neurons tested in N = 14 mice, with qualitatively similar spiking observed with train stimulation in n = 5 VGAT-neurons from N = 3 mice recorded in cell-attached mode (Figure 7A. Median fraction of trials with spikes: 0.63). Moreover, the initial EPSPs which depolarized VGAT-neurons towards spike threshold closely followed each light flash in the train (Figure 7B). These data suggest that monosynaptic descending inputs suffice to drive spiking independently of ascending activity.

In VGAT+ neurons, repetitive corticofugal stimulation drove spikes in n = 21/45 neurons tested in N = 21 mice (47%; Figure 7C. Median fraction of trials with spikes: 0.24). These values likely represent a lower bound, as recordings were generally conducted while hyperpolarizing neurons with negative bias current to limit tonic firing and observe the underlying synaptic depolarization. Interestingly, trace-by-trace inspection of synaptic activity during train stimulation suggested that unlike the monosynaptic EPSPs seen in VGAT-neurons, the EPSPs driving spikes in VGAT+ neurons often bore the hallmarks of a polysynaptic origin. Individual events had a variable and delayed onset relative to each light flash, sometimes occurring as barrages of summating events (Figure 7D). These data suggest that monosynaptic corticofugal activity drive spikes in a subset of VGAT-(e.g., glutamatergic) neurons, thereby leading to feedforward excitation in VGAT+ neurons.

To further determine the prevalence of mono-and di-synaptic activity during corticofugal stimulation, we compared the properties of *sub-threshold* EPSPs in VGAT-and VGAT+ neurons during corticofugal train stimulation (25 flashes at 50 Hz). As in Figure 7, train stimuli typically generated short latency EPSPs in VGAT-neurons that were tightly locked to the individual light flashes and displayed temporal summation (Figure 8A,B; n = 28 neurons from N = 14 mice). Long-latency and temporally jittered EPSPs were rarely observed in these recordings (n = 2/28 neurons tested, or 7%; data not shown), suggesting that corticofugal activity recruits minimal polysynaptic activity in VGAT-neurons. By contrast, over half of VGAT+ neurons tested displayed a strikingly distinct profile (n = 25/46 neurons, or 54%; Figure 8C,D): Even under conditions where the first train stimulus resulted in weak short-latency EPSPs, subsequent light flashes resulted in jittered barrages of large amplitude EPSPs with delayed onset latencies (Figure 8D). We interpret the initial short-latency EPSP as reflecting the monosynaptic input from corticofugal synapses (Figure 8D, arrowheads), while the later, more temporally-jittered EPSPs reflect disynaptic excitation (Figure 8D, asterisks) from local glutamatergic neurons driven to spike by monosynaptic corticofugal inputs. In the other population of VGAT+ neurons tested (21/46 neurons, or 46%), we observed only short-latency, monosynaptic EPSPs (Figure 8E,F). As such, the overall probability of observing recurrent excitation during corticofugal train stimulation was significantly higher in VGAT+ compared to VGAT-neurons (χ^2^= 14.761, p = 0.0001). These EPSP timing differences were further apparent when averaging the peak normalized membrane potential waveform following each stimulus cycle in the 50 Hz train: The cycle average waveform for the VGAT+ neuron with long-latency EPSPs clearly shows two distinct peaks which presumably reflect mono-and poly-synaptic excitation during repetitive corticofugal train stimulation (Figure 8G, magenta). By contrast, the other two example recordings show a single short-latency peak, reflecting monosynaptic inputs from corticofugal synapses (Figure 8G, black and green).

To test if EPSP latency differences between the three groups were statistically significant, we generated summary histograms (0.4 ms bin-widths) for EPSP latencies following each light flash in the train, and normalized the event counts in each bin to the total number of observations in each recording. The mean histograms from VGAT-neurons and VGAT+ neurons that lacked polysynaptic EPSPs showed a single peak 3-5 ms following light onset (Figure 8H; black and green, respectively). By contrast, the histogram from VGAT+ neurons with polysynaptic EPSPs was clearly bi-modal, showing an initial peak near 3-5 ms followed by a second peak near the 7-8 ms latency bins (Figure 8H, magenta). A two-way, repeated measures ANOVA comparing latency histograms revealed main effects of latency bin (F(3.832, 272.1) = 41.69, p < 0.0001), neuron type (F(2, 71) = 37.86, p < 0.0001), and a latency bin x neuron type interaction (F(98, 3479) = 6.736, p < 0.0001).

The bi-modal EPSP distribution seen in group data was not an artifact of averaging histograms from two sub-populations of neurons with short-and long-latency peaks. Rather, the short-and long-latency peaks were also observed in EPSP latency histograms from individual recordings of VGAT+ neurons (Figure 9A-C). Importantly, this bi-modal timing further rules out alternative hypotheses such as asynchronous release to explain the jittered and long-latency EPSP barrages: A bi-modal latency distribution is expected from a scenario where direct monosynaptic excitation is quickly followed by recurrent polysynaptic inputs (Molnár et al., 2008; Pita-Almenar et al., 2014; Hemberger et al., 2019). By contrast, asynchronous release follows an exponential distribution after each presynaptic action potential (Best and Regehr, 2009; Turecek and Regehr, 2018). Altogether, our results show that repetitive corticofugal activity drives feedforward excitation in a major fraction of dorsal IC VGAT+ neurons, thereby providing a circuit mechanism to amplify corticofugal signals.

**Figure 9:**
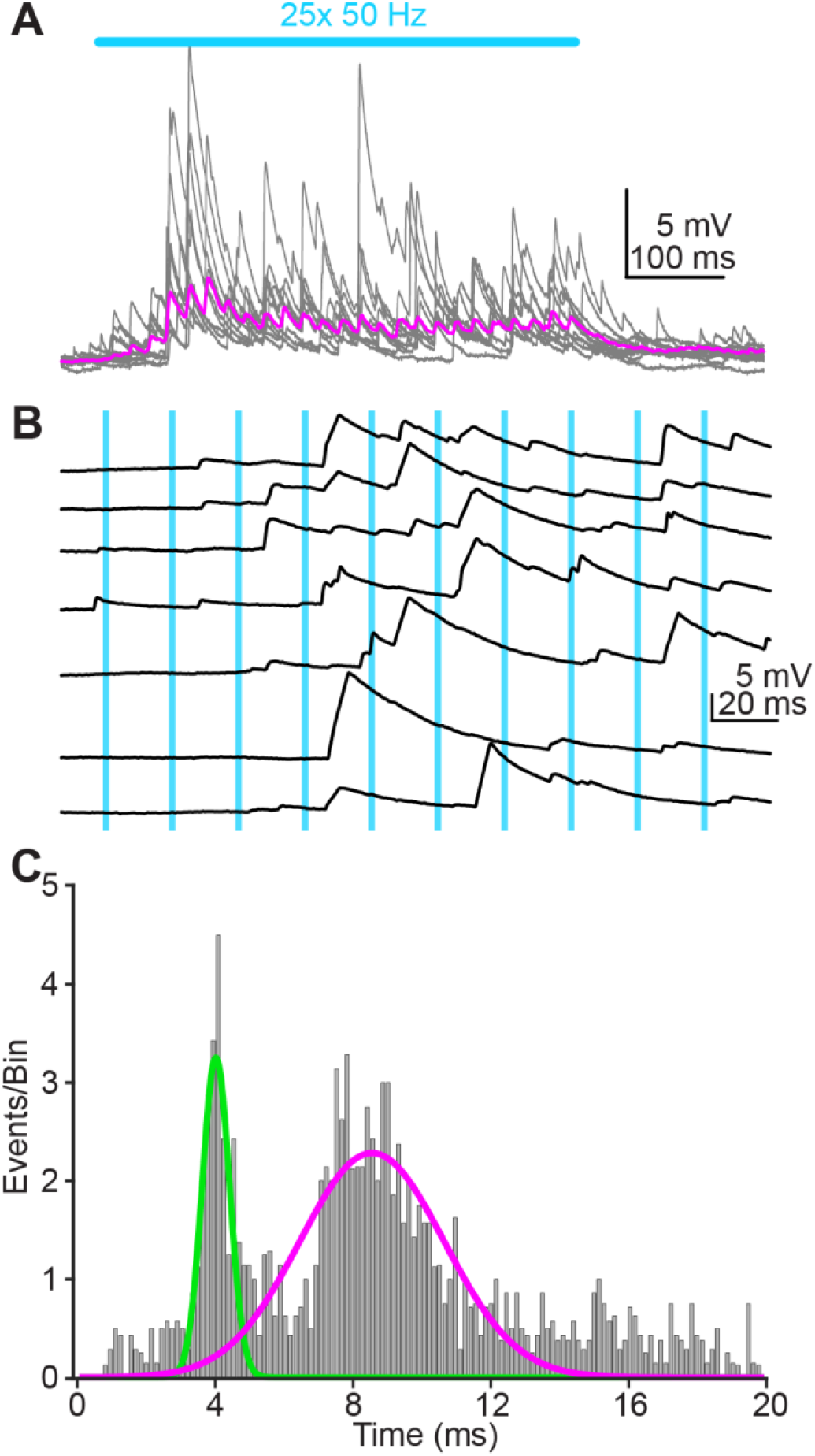
Additional examples of polysynaptic EPSPs in a VGAT+ neuron. **A)** Example single trials (gray) and average (magenta) of synaptic barrages during 50 Hz corticofugal stimulation in a VGAT+ neuron. Blue bar denotes optogenetic stimulation. **B)** A subset of the single trials from panel A are expanded to highlight EPSPs evoked during the first 10 light pulses. Blue lines indicate onset of individual light flashes. **C)** Latency histogram for EPSPs occurring following each light flash in the train. X-axis is limited to 0-20 ms, reflecting the duty cycle of a 50 Hz stimulus train. Of note, the latency histogram shows two distinct peaks, suggesting that EPSPs arise from both mono-and poly-synaptic inputs. Green and magenta curves are fits from a two-term Gaussian model.

**Figure 10:**
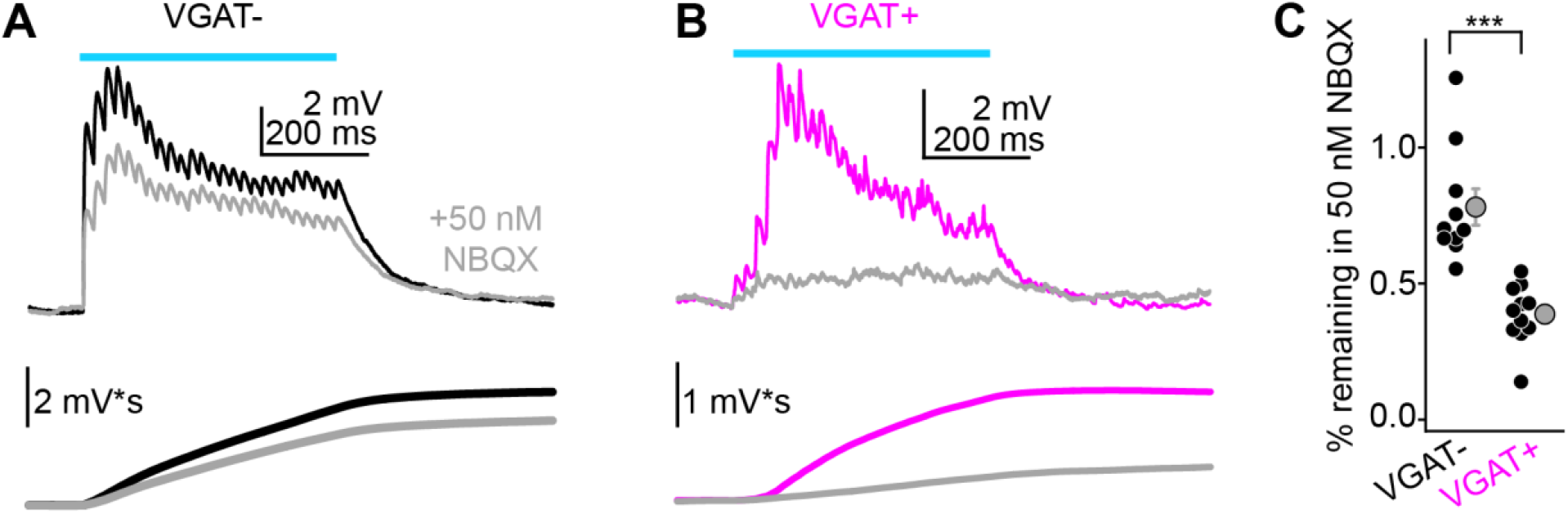
Differential block of “corticofugal” excitation in VGAT-and VGAT+ neurons by 50 nM NBQX. **A,B)** Example average train EPSPs from a shell IC VGAT-(A) or VGAT+ (B) neuron before (black and magenta) and after (gray) bath application of 50 nM NBQX. Lower traces are the cumulative integral of the EPSP waveform. **C)** Summary data showing the fraction remaining in 50 nM NBQX for the cumulative integral of the EPSP waveform. Black and gray symbols are individual experiments and mean ± SEM, respectively. *** p<0.0001.

**Figure 11:**
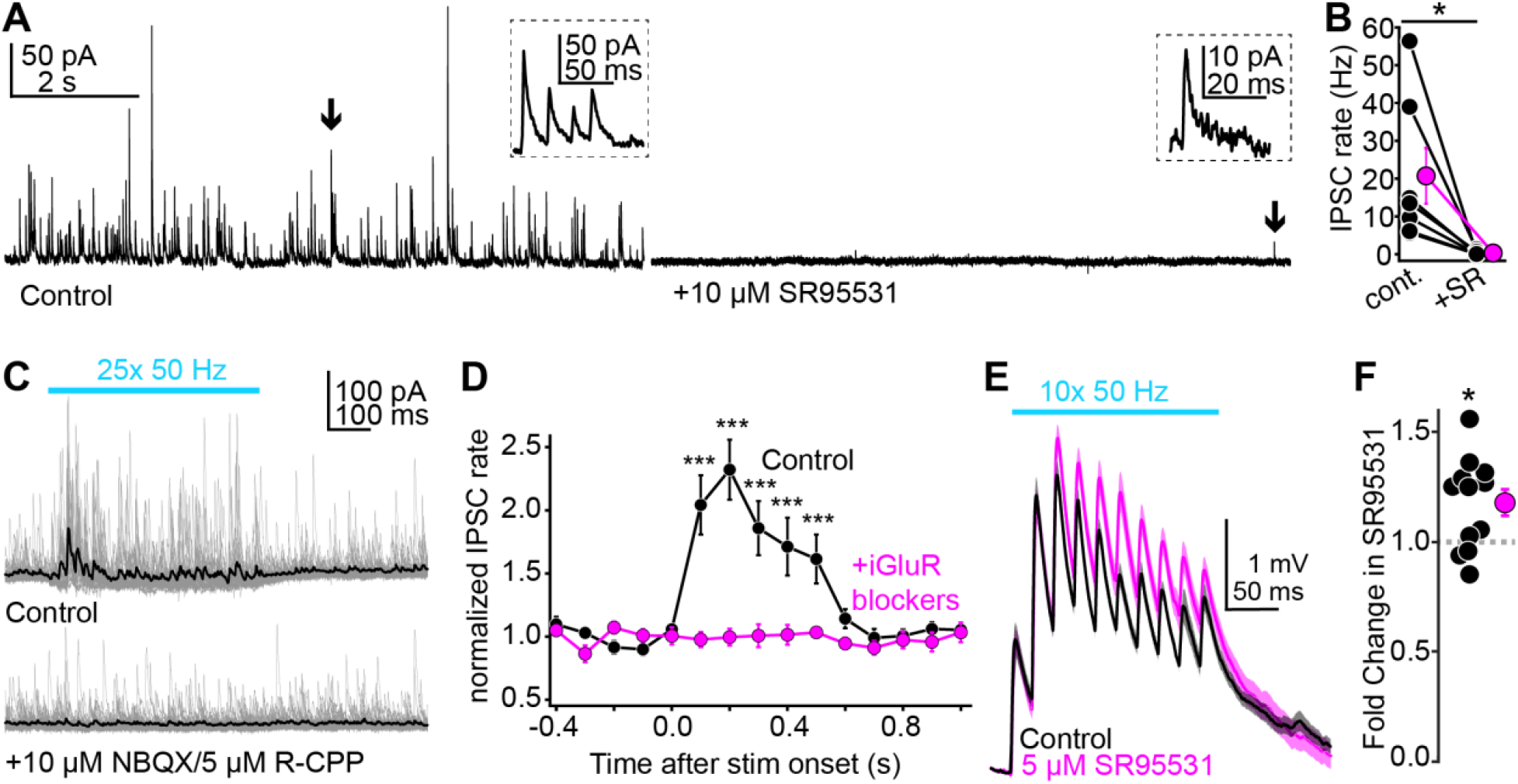
Corticofugal activity generates polysynaptic inhibition in the dorsal IC. **A)** Left: A dorso-medial IC neuron voltage-clamped at +8 mV in control conditions. Of note are the spontaneous IPSCs often occurring in bursts, suggesting they are mediated by action potentials in local GABA neurons. Arrow denotes one such IPSC burst shown at a faster time base in the inset. Right: Bath application of the GABAA receptor antagonist SR95531 (10 µM) blocks the majority of spontaneous IPSCs. Although quite rare, a few events nevertheless remained in the presence of SR95531 (arrow and inset; median IPSC rate in SR95531: one event per 17.95 seconds). **B)** Summary data for n = 7 neurons showing that SR95531 profoundly reduces spontaneous IPSC rates. Thus, inhibitory transmission in dorsal IC neurons is mostly GABAergic. **C)** Example recording from a dorsal IC neuron before and after bath application of glutamate receptor blockers. Gray traces are overlays of individual trials, black is average. Of note is that the powerful increase in IPSC rate during corticofugal stimulation (blue bar) is abolished by blocking glutamate receptors. **D)** Normalized IPSC rate is binned every 100 ms for statistical comparisons. Asterisks denote significance in Sidak’s post-hoc tests comparing event rates across control and drug bins. Of note is that significance is limited to timepoints during optical stimulation of corticofugal axons. **E)** Average traces showing temporal integration of corticofugal EPSPs in current-clamp before (black) and after (magenta) bath application of the GABAA receptor antagonist SR95531. Shading is ± SEM. **F)** Summary data, Values are the ratio of voltage integrals in drug and control conditions. * p <0.05, *** p<0.001.

### Dampening network activity differentially reduces corticofugal depolarizations in VGAT-and VGAT+ neurons showing long-latency EPSPs

If the asynchronous barrages in VGAT+ neurons reflect polysynaptic EPSPs from presynaptic glutamatergic neurons with local axons, the corticofugal depolarization in this major fraction of VGAT+ neurons should be uniquely sensitive to experimental manipulations that reduce feedforward spiking of VGAT-neurons. By contrast, the mostly monosynaptic corticofugal EPSPs in VGAT-neurons should be comparatively less reduced by the same manipulation. To test this prediction, we asked if a sub-saturating concentration of the AMPA receptor antagonist NBQX (50 nM) causes a greater reduction of corticofugal activity in VGAT+ neurons with long-latency EPSPs compared to VGAT-neurons, which largely lacked long-latency EPSPs. This concentration of NBQX blocks 20-50% of the AMPA receptor charge transfer (Randle et al., 1992; Diamond and Jahr, 1997) and will decrease the probability that IC glutamatergic neurons fire spikes during corticofugal stimulation, thus non-linearly reducing polysynaptic EPSPs in VGAT+ neurons. Furthermore, NBQX is a high affinity blocker, and diffusion of glutamate out of the synapse (Clements et al., 1992) is faster than the antagonist unbinding rate; the degree of block of by NBQX is thus independent of any hypothetical differences in presynaptic release probability or cleft glutamate concentration between VGAT-and VGAT+ neurons (Christie and Jahr, 2006). Rather, a greater degree of block in VGAT+ neurons should reflect a non-linear reduction in polysynaptic activity when partial AMPA receptor blockade reduces corticofugal-evoked spiking of VGAT-neurons.

Accordingly, 50 nM NBQX reduced the cumulative voltage integral during stimulus trains by 22 ± 7% in VGAT-neurons (Figure 10A,C; n = 10 neurons from N = 8 mice). By contrast, 50 nM NBQX caused a significantly greater reduction in VGAT+ neurons showing long-latency and asynchronous EPSPs (61 ± 3% reduction, Figure 10B,C; n = 11 neurons from N = 8 mice; p < 0.0001 compared to glutamate neurons, paired t-test). These data argue that while auditory cortical activity strongly depolarizes VGAT+ neurons (Figure 6 and 7), much of the underlying depolarization originates from local excitatory inputs rather than monosynaptic connections from corticofugal axons.

### Corticofugal activity generates polysynaptic inhibition

The net depolarization during corticofugal train stimulation was often large enough to drive spikes in VGAT+ neurons (Figure 7C,D). If VGAT+ neurons project their axons locally, corticofugal activity could thus suffice to generate feedforward inhibition in the dorsal IC. To test this hypothesis, we injected wild-type mice with a Chronos virus in auditory cortex and voltage-clamped dorsal IC neuron somata near the reversal potential for excitation (∼ +5 to +10 mV) to record inhibitory postsynaptic currents (IPSCs). Spontaneous outward currents under these conditions overwhelmingly reflected activity at GABAergic synapses, as they were almost completely abolished by the GABAA receptor antagonist SR95531 (5-10 µM; Figure 11A,B. Median IPSC rate: 13.5 vs. 0.0557 Hz in control and SR95531, p = 0.0156, sign-rank test, n = 7 neurons from N = 3 mice). The few remaining events in SR95531 may reflect glycinergic IPSCs from long range axons originating in the ventral nucleus of the lateral lemniscus (Moore & Trussell, 2017).

Stimulating corticofugal axons with trains of blue light flashes (25x, 50 Hz) resulted in a powerful, ∼2-3 fold increase in IPSC rate relative to baseline (Figure 11C), as revealed by a main effect of time in a two-way, repeated measures ANOVA (Figure 11D. F(14,126) = 11.7, p < 0.0001, n = 10 neurons from N = 5 mice). Importantly, the IPSC rate increase was abolished by bath application of glutamate receptor antagonists NBQX (10 µM) and/or R-CPP (5 µM), indicating that it was mediated by polysynaptic activation of IC GABA neurons (Figure 9C,D. Timepoint x drug interaction: F(14,126) = 13.38, p < 0.0001). Functionally, this feedforward inhibition controlled the temporal integration of corticofugal excitation: Blocking GABAA receptors significantly increased corticofugal EPSP summation during repetitive stimulation in current clamp, measured as a significant increase in the voltage integral following bath application of 5-10 µM SR55931 (Figure 11E,F. Fractional change relative to control: 1.18 ± 0.06, n = 12 neurons from N = 11 mice. p = 0.013, one sample t-test). Interestingly, we also noted a minor, but statistically significant increase in membrane resistance following SR95531 (386 ± 61 vs. 397 ± 62 MΩ in control and SR95531, respectively. Fractional change: 1.04 ± 0.014, p = 0.0204, one sample t-test). Although this observation argues that the high rates of spontaneous IPSCs from the local circuit contribute a certain level of effectively tonic inhibition (e.g., Figure 11A; See also Brickley et al., 1996, 2001), this small change in membrane resistance in SR95531 is unlikely to account for our observed effect on EPSP summation: The relative increase in EPSP integral was more than 4-fold larger than the change in membrane resistance, and there was no significant pairwise correlation between the magnitude of EPSP enhancement and change in membrane resistance after SR95531 application (Pearson’s coefficient ± standard deviation from 100 bootstrapped iterations: -0.25 ± 0.36, p = 0.43). Thus, corticofugal transmission generates intra-collicular inhibition that regulates how IC neurons integrate descending signals over time.

## Discussion

Although decades of *in vivo* physiology show that auditory cortical activity inhibits IC sound responses, the underlying synaptic mechanisms have been difficult to pin down. Indeed, anatomy data suggest that auditory cortical axons preferentially target glutamate, rather than GABA neurons in the IC (Nakamoto et al., 2013; Chen et al., 2018). This anatomical motif is in striking contrast to the often studied pyramidal neuron-based microcircuits of sensory cortices and hippocampus, where long-range excitation powerfully drives feedforward inhibition to sharpen temporal fidelity of afferent signals: Glutamatergic synapses onto GABAergic interneurons in these circuits are typically more numerous, have higher synaptic conductance, and/or higher release probability than onto pyramidal projection neurons (Acsády et al., 1998; Lawrence et al., 2004; Gabernet et al., 2005; Cruikshank et al., 2007). At first glance, the anatomical constraints of the auditory cortico-collicular pathway suggest that descending inhibition of the IC is unlikely to operate via a classical feedforward recruitment of locally projecting IC GABA neurons. However, our experiments reveal cellular and circuit mechanisms that enable the auditory cortex to trigger intra-collicular inhibition in absence of a dense innervation onto IC GABA neurons. Consistent with the anatomical data (Nakamoto et al., 2013; Chen et al., 2018), corticofugal synapses tended to generate smaller EPSPs in VGAT+ compared to VGAT-neurons. Nevertheless, two cellular mechanisms enabled corticofugal activity to effectively increase firing rates of IC GABA neurons. First, approximately half of dorsal IC GABA neurons encountered in our study fired tonically, similar to findings in the NPY-positive GABA neurons of the central IC (Silveira et al., 2020). Thus, even apparently weak descending EPSPs sufficed to increase the basal firing rates of GABA neurons. Second, repetitive firing of corticofugal axons drives robust polysynaptic excitation in IC GABA neurons due to local excitatory circuitry, thereby orchestrating a multisynaptic cascade leading to intra-collicular inhibition. An important caveat is that our results cannot rule out if in addition to intra-collicular circuit mechanisms, naturalistic patterns of auditory cortical activity recruit other long-range GABAergic projections to inhibit IC neurons. Indeed, corticofugal synapses may contact neurons in the dorsal nucleus of the lateral lemniscus (Budinger et al., 2000), and neurons of the nucleus sagulum which project to the IC and may be GABAergic (Henkel and Shneiderman, 1988; Beneyto et al., 1998). However, the corticofugal projection to sub-collicular targets is much sparser than the auditory cortico-collicular pathway (Doucet et al., 2003; Coomes and Schofield, 2004; Schofield et al., 2006). Thus, reliable recruitment of long-range inhibition may require particularly strong corticofugal synapses, or alternatively rely on amplificatory mechanisms such as those described herein. Future studies, potentially employing synapse-specific silencing approaches (Copits et al., 2021), are needed to disentangle the relative contribution of local and long-range inhibition to corticofugal suppression of IC activity.

We did not observe direct monosynaptic GABAergic inputs from auditory cortex to the dorsal IC, as recently suggested by anatomy data (Bertero et al., 2021). *In vitro*, the IPSC barrages recruited during auditory cortex stimulation were entirely blocked by glutamate receptor antagonists, indicating the necessity for intermediary, excitatory synapses onto IC GABA neurons with locally projecting axons. This result is perhaps surprising, as our stimulation paradigm was quite strong (25 light flashes at 50 Hz) and likely sufficed to activate descending synapses with low release probability.

Furthermore, Bertero et al. showed that long-range GABAergic axons from auditory cortex terminate in the dorsomedial IC shell where our recordings were obtained. However, an absence of evidence is not conclusive evidence of absence: This newly discovered pathway may instead operate via non-canonical mechanisms such as extrasynaptic diffusion of GABA, or via release of the neuropeptide VIP. These synaptic properties would be difficult to quantify with our whole-cell patch-clamp recordings. In addition, selective viral tropism may have hindered the expression of our optogenetic construct in auditory cortical GABAergic neurons; extensive follow-up experiments are necessary to clarify these issues. Rather, our results show that intra-collicular mechanisms suffice to explain at least some of the auditory cortex’s inhibitory actions on IC sound responses.

Previous studies showed that the majority of IC GABA neurons display an HCN channel dependent, depolarizing “sag” during negative current injections. By contrast, we only observed similar membrane properties in a subset of our recordings from VGAT+ IC neurons (Ono et al., 2005). These discrepancies may reflect age-or sub-division dependent biases in the membrane properties of IC neurons. Indeed, Ono et al. recorded from GABA neurons located across the central, dorso-medial, and lateral subdivisions and their experiments were restricted to animals between ∼2-4 weeks of age. By contrast, we selectively targeted neurons in the dorso-medial IC in 5-9 week old mice, which may explain the discrepancies between our work and that of Ono et al.

Prior experiments which stimulated auditory cortex *in vivo* did not report recurrent excitation in IC neurons. However, this absence is perhaps not surprising, as we find that polysynaptic EPSPs are primarily restricted to VGAT+ neurons. Given that GABAergic neurons represent ∼25% of the total IC neuron population, the previous studies employing “blind” extracellular recordings in the IC likely under-sampled from GABAergic neurons. Furthermore, unambiguous identification of polysynaptic EPSPs requires intracellular recordings. To our knowledge, only three prior studies performed intracellular recordings from IC neurons during cortical stimulation (Mitani et al., 1983; Qi et al., 2020; Oberle et al., 2022), and none of these recordings were targeted to GABAergic neurons. Rather, the fact that corticofugal activity preferentially triggers polysynaptic EPSPs in VGAT+ neurons suggests that local connectivity in the IC is not random, but rather follows specific wiring rules. Accordingly, the somata of “large GABAergic” neurons in the IC are densely innervated by VGluT2+ boutons (Ito et al., 2009; Ito and Oliver, 2014). These somatic boutons may originate from neighboring excitatory IC neurons, such that large GABAergic neurons may be preferentially activated by local activity. Further studies, perhaps employing cell-type specific markers for sub-types of IC GABA neurons, will be required to fully determine how local and long-range inputs activate distinct IC cell-types.

Functionally, the polysynaptic activity driven by descending transmission could enable a myriad of “high-level” computations in the shell IC. For example, previous work in piriform cortex shows that recurrent excitatory synapses from principal cells onto interneurons trigger feedback inhibition and truncates the duration of principal cell responses to incoming activity from the olfactory bulb. Although odorant signals from the bulb’s lateral olfactory tract are graded with respect to odorant concentration, the output of piriform cortex principal cells is consequently rendered concentration invariant by this feedback inhibition which powerfully limits spiking to odor onset (Bolding and Franks, 2018). Our findings reveal a conceptually similar microcircuit architecture in the dorsal IC. Corticofugal projections, which potentially transmit behaviorally relevant information, could thus enhance IC glutamate neurons’ ability to trigger feedback inhibition and promote a level-invariant representation of behaviorally relevant sounds in the IC. Such a computation may be particularly useful in creating invariant neural responses necessary for sound localization in complex, reverberant environments (Hartmann, 1983; Rakerd and Hartmann, 1986). Indeed, the onset spiking response of IC neurons represents directional signals that are less corrupted by reverberant reflections and inter-aural decorrelation than later, sustained responses (Devore et al., 2009; but see Barzelay et al., 2023). It is not unreasonable to assume that onset spiking in glutamate neurons could recruit feedback inhibition that minimizes sustained activity during the reverberant tail of sounds, and that this mechanism is enhanced by “top-down” cortical signals during periods of increased vigilance and attention. However, extensive future studies are required to elucidate the behaviorally relevant conditions that trigger descending inhibitory control, and to understand how corticofugal projections might flexibly switch between amplifying and inhibiting tectal sound responses.

## Acknowledgments

Funding was provided by the Whitehall Foundation, Hearing Health Foundation, and NIH/NIDCD R01DC019090 to PFA, as well as T32DC005356 and T32DC000011 to HMO. We thank Drs. Stephanie Gantz and Michael Roberts for critical comments on the manuscript, and Audrey Drotos for preliminary analyses of EPSP timing in VGAT+ neurons.

